# Therapeutic Potential of the Novel Monoclonal Antibody 7-4G against HBsAg in a Mouse Model of Chronic Hepatitis B Virus Infection

**DOI:** 10.64898/2026.07.24.740663

**Authors:** Maidina Abuduwaili, Grace Naswa Makokha, Malihe Naderi, Hiroyuki Sakai, Akima Yamamoto, Takemasa Sakaguchi, Takehisa Matsumoto, Mikako Shirouzu, Kyoizumi Seishi, Ayako Yamamoto, Yoshiko Kubo, Miyuki Matsushita, Yoko Inoue, De Shankhajit, Makoto Hijikata, Kazuaki Chayama

**Author notes:** These authors contributed equally to this work (Maidina Abuduwaili, Grace Naswa Makokha).

## Abstract

Chronic hepatitis B virus (HBV) infection remains a major global health challenge because current antiviral therapies rarely achieve complete viral clearance. We evaluated the antiviral activity of a newly developed monoclonal antibody, 7-4G mAb, directed against the hepatitis B surface antigen (HBsAg). The antibody showed specific binding to the p27 component of HBsAg and efficiently removed HBV-related particles from patient serum in vitro. Significant HBV neutralizing activity of 7-4G mAb was observed in primary human hepatocytes and in humanized mouse models. In persistently HBV-infected PXB mice, a single administration of 7-4G mAb, either alone or in combination with another monoclonal antibody, reduced serum HBsAg levels and HBV DNA levels by two orders magnitude for two weeks after the treatment. In half of the treated mice, suppression of both viral markers was maintained through the study period, whereas rebound occurred within three to four weeks in the remaining animals. In addition, 7-4G mAb enhanced the HBV DNA-lowering effect of entecavir, a known anti-HBV drug. These findings support the further development of 7-4G mAb as part of an innovative HBV treatment strategy, addressing limitations of existing therapies and advancing toward a more effective HBV management solution.

## Introduction

Hepatitis B virus (HBV) infection affects approximately 240 million people worldwide and remains a major global health burden (1). Chronic infection is particularly common when exposure occurs during infancy, where immune tolerance allows persistent viral replication and continuous production of hepatitis B surface antigen (HBsAg) (2). High circulating levels of HBsAg - largely derived from excessive secretion of subviral particles (SVPs) - are a defining feature of chronic hepatitis B (CHB) and contribute to impaired antiviral immunity (3). Because loss of HBsAg is the central requirement for achieving a functional cure, strategies capable of reducing circulating HBsAg and SVPs are urgently needed. Current antiviral therapies, including nucleos(t)ide reverse-transcriptase inhibitors (NRTIs) and PEGylated interferon-α (PEG-IFNα), suppress HBV replication but rarely eliminate HBsAg (4). NRTIs efficiently reduce serum HBV DNA yet have minimal impact on HBsAg levels, resulting in lifelong therapy for most patients. PEG-IFNα can inhibit transcriptional activity of covalently closed circular DNA (cccDNA) (5,6), but its use is limited by substantial side effects and restricted patient eligibility (7). Consequently, the rate of functional cure - defined as complete HBsAg loss with or without seroconversion - remains extremely low, typically 1-10% after years of treatment (8,9). These limitations highlight the need for therapeutic approaches that directly target HBsAg and SVPs, rather than viral replication alone.

HBsAg - targeting antibodies represent a rational strategy to overcome this barrier. SVPs are produced at quantities far exceeding infectious virions and act as decoys that blunt antiviral immune responses (3). Henceforth, their removal could restore immune recognition of HBV and facilitate HBsAg clearance. Polyclonal hepatitis B immunoglobulin (HBIG) demonstrates that antibody-mediated neutralization can be effective (10), but its clinical use is constrained by high cost, limited donor-derived supply, and biosafety concerns associated with blood products (11). In contrast, monoclonal antibodies (mAbs) offer consistent quality, scalable production, and improved safety, making them attractive candidates for HBV immunotherapy. Currently, there are nearly forty mAbs in clinical development worldwide, targeting more than a dozen infectious diseases including human immunodeficiency virus (HIV), anthrax, Ebola, hepatitis, and influenza (12). Moreover, in recent years, human mAb therapeutics have been attracting attention again because of the COVID-19 pandemic (13). On the other hand, although several mAbs targeting HBV have been described, most focus on the preS1 region of the large HBsAg protein, which mediates viral entry via NTCP (14–18). Far fewer antibodies selectively recognize conformational epitopes on the small HBsAg proteins (p27/p21), despite their abundance on SVPs and Dane particles. Antibodies directed against native conformational epitopes of small HBsAg may be particularly effective in clearing circulating antigen and neutralizing infection across HBV genotypes.

In this study, we developed and characterized a novel monoclonal antibody, 7-4G, which specifically binds a conformational epitope on the native p27 component of HBsAg. We evaluated its antiviral activity in vitro using primary human hepatocytes and in vivo using humanized liver chimeric PXB mice. Furthermore, we assessed its therapeutic potential in chronic HBV infection and examined its additive effects when combined with the antiviral agent entecavir (ETV). Our findings support the development of 7-4G mAb as a promising component of next-generation HBV treatment strategies aimed at achieving functional cure.

## Materials and Methods

### HBV infectious serum

The sera from patients infected with HBV were collected at the Hiroshima university hospital and Hiroshima institute of life sciences and used for the studies in accordance with the guidelines of the local ethics committees of Hiroshima university and the Hiroshima institute of life sciences.

### Experimental models

PXB mice are chimeric mice with humanized livers and were purchased from Phoenix Bio co. Ltd. (Hiroshima, Japan). The mice were established from a host strain that has a transgene containing albumin promoter/enhancer and urokinase-type plasminogen activator cDNA and has a SCID background (cDNA-uPA/SCID) (19). Primary hepatocytes freshly isolated from the PXB mice (PXB cells) were also separately purchased from Phoenix Bio co. Ltd. The cells were cultured on collagen-coated 24-well plates (BioCoat, Corning, NY, USA) in dHCGM, a culture medium also supplied by Phoenix Bio co. Ltd, Hiroshima, Japan. The animal experimental protocols were in strict adherence to the ethical guidelines of the Declaration of Helsinki, and were approved by the Hiroshima university ethical committee in accordance with the guidelines of the local committee for animal experiments by the committee for recombinant DNA experiments and animal experiments of Hiroshima university as well as the Hiroshima institute for life sciences.

### Development of mAbs against HBsAg

The mAb targeting HBsAg of HBV genotype C in the natural form, was developed using the HBV particles isolated from the patient’s serum with ultracentrifugation as an antigen via standard hybridoma technology by the Institute of Immunology (ITM Co., Ltd, Nagano, Japan). The anti-HBsAg activity in the culture medium of the hybridoma were evaluated with anti-HBsAg enzyme-linked immune-solvent assay (ELISA). The anti-HBsAg positive hybridomas were selected and used to produce the mAbs.

### Validation of specificity of 7-4G mAb for the antigen structure

Denatured antigen was prepared by incubating 5 μL (5 μg) of 1 mg/mL HBsAg (Kamakura Techno science. Inc., Kanagawa, Japan) in 50 mM Tris-HCl, 2% sodium dodecyl sulfate (SDS), 5% 2-mercaptoethanol (2ME) (pH 7.0) at 95°C for 3 min, followed by dilution to make a 1 μg/mL solution. The denatured antigen solution and the native form of HBsAg, also diluted in PBS to 1 μg/mL was added to 96 well ELISA plates (Nacalai Tesque, Kyoto, Japan) at 100 uL/well, and incubated overnight at 4°C. After the incubation and blocking with Block one solution (Nacalai Tesque, Kyoto, Japan), 100 μL/well of purified antibody in eight concentrations starting from 1 μg/mL at 3-fold step dilution was added to the wells. Rabbit Anti-mouse IgG Fc fragment Horseradish Peroxidase (HRP)-Conjugated (Thermo Fisher Scientific, Tokyo, Japan) was used for detection. HRP was reacted with tetramethylbenzidine (TMB) Substrate (Nacalai Tesque, Kyoto, Japan) and the reaction stopped using the TMB Substrate stop Solution (Nacalai Tesque, Kyoto, Japan). Absorbance at 450 nm was detected using the Multiskan SkyHigh Microplate Spectrophotometer (Thermo Fisher Scientific, Tokyo, Japan).

### Identification of HBV particles bound with 7-4G mAb

Serum from HBV-infected patients (0.5 mL) was mixed with purified 7-4G mAb (0.25 mL, 0.25 mg) and incubated on ice for several hours. For the negative control, PBS was used instead of the antibody. The resulting mixture was layered onto a 20%–60% (w/v) sucrose density gradient in PBS and ultracentrifuged at 25,000 rpm and 4°C for 16 h using a Beckman Optima L-90K ultracentrifuge and an SW 41 Ti rotor. Following centrifugation, the gradient was fractionated into 0.5 mL aliquots using an Auto Densi-Flow fractionator (Labconco, Kansas City, MO, USA). For each fraction, sucrose density, HBsAg levels, and HBV DNA levels were measured.

### Identification of HBsAg polypeptides recognized by 7-4G

Fifty μg of control mouse IgG or 7-4G mAb was adsorbed to Protein G Sepharose 4 Fast Flow (Cytiva, Marlborough, Massachusetts, USA) resuspended in 100 μL of HBS buffer (20 mM HEPES pH 7.5, 150 mM NaCl) containing 0.1% Tween 20 for 2 hours at room temperature. After washing with HBS buffer containing 0.1% Tween 20, the beads were incubated with 170 μg of crude HBV particles isolated from the patient’s serum with ultracentrifugation for 3 hours at room temperature. After incubation, each set of beads was washed five times with HBS buffer containing 0.1% Tween 20. An equal volume of 2x Sample Loading buffer (containing 2ME) was added to the washed beads and heated at 95°C for 5 minutes. After brief centrifugation, the supernatant was separated by SDS-polyacrylamide electrophoresis (PAGE) followed by staining with Coomassie Brilliant Blue (CBB) or silver stained with Sil-Best Stain One (Nacalai Tesque, Kyoto, Japan).

### Quantitation of HBV markers

Quantitative analyses of HBsAg, HBeAg and anti-HBs antibodies (HBsAb) in the culture medium of HBV infected PXB cells and sera from HBV infected mice were outsourced to SRL Laboratory (SRL, inc. Tokyo, Japan). Determination of the HBV DNA level in the sera of HBV infected mice was done by quantitative polymerase chain reaction (qPCR) using HBV specific primers as described previously (20).

### In vitro neutralization assay

HBV infection was performed as we previously described (21). Briefly, PXB cells were seeded onto collagen coated 24-well plates at a concentration of 1 × 10⁵ cells per well and the serum of a patient with chronic HBV infection of genotype C,-with high virus load (equal or higher than 10^9^ copies/mL) was utilized for infection. Cells were exposed to the infectious serum at a genome equivalent (GEq) of 100, in the presence of varying doses of mAbs at 0.01, 0.1, and 1 μg/mL. At thirteen days after the inoculation, the levels of HBeAg and HBsAg in the collected culture supernatants were measured as described above.

### In vivo neutralization assay

PXB mice were administered with no antibody, anti-FLAG antibody at 0.5 mg/mouse (negative control), the anti-preS1 antibody 3-10 (3-10 mAb), previously established (22) at 1 mg/mouse, or the 7-4G mAb at 0.5 mg/mouse via intravenous tail vein injection. For the combination treatment, 0.33 mg of 3-10 mAb and 0.66 mg of 7-4G mAb were administered using the same regimen. One day after the inoculation, these mice were injected with serum of patients chronically infected with HBV genotype C including 10^3^ GEq HBV per mouse via the tail vein, subsequently. The blood samples of the mice were collected on a weekly basis for up to seven weeks from which the serum levels of HBsAg and HBV DNA were measured using ELISA and qPCR, respectively, as described above. For examination of the neutralization effect of 7-4G mAb on other HBV genotypes, HBV inocula of genotypes A, B, and D which consisted of sera from virus infected PXB mice was utilized for infection of PXB mice, subsequent to pretreatment with 50 mg/kg/mouse of 7-4G mAb. Sera produced in mice was used for these experiments due to the limited availability of patient sera for HBV infection experiments, however, the original inocula used to infect the mice for amplification was obtained from HBV infected patients at Hiroshima University, after an informed consent.

### The effect of 7-4G mAb on chronic HBV infection in vivo

PXB mice were injected with the serum of patients chronically infected with HBV genotype C including 10^3^ GEq HBV per mouse via the tail vein. The blood samples of the mice were collected on a weekly basis, and the serum levels of HBV DNA were monitored for up to the twenty third week after inoculation. Fourteen or fifteen weeks after the inoculation, when HBV genome titers in the PXB mice seemed to reach the plateau levels, three mice in each group were injected with 0.5 mg/mouse of anti-FLAG control antibody, 7-4G mAb (1 mg/mouse), 3-10 mAb (2 mg/mouse), or a combination of both antibodies (1 mg of 7-4G mAb and 2 mg of 3-10 mAb /mouse). Thereafter, the serum levels of HBsAg, and HBsAb were monitored weekly until week twenty-three, as the case of HBV DNA.

### Combination treatment of entecavir and 7-4G mAb

PXB mice were inoculated with serum of patients chronically infected with HBV genotype C as described above for chronic infection. Nine weeks after the inoculation, the mice were injected with the anti-HBV agent, entecavir (ETV), at a dose of 0.2 mg/kg/day for up to fifteen weeks (ETV alone group). For the ETV and 7-4G mAb combination group, the mice were treated with ETV from week nine as above, then at the thirteenth week post inoculation, they were injected with 7-4G mAb (1 mg/mouse). Blood samples of the mice were collected on a weekly basis, and the serum levels of HBV DNA were monitored for up to the eighteenth week after inoculation.

## Statistical analysis

Where applicable, data are presented as means ± standard errors of the means (SEM).

## Results

### 1. Isolation of hybridoma producing mAbs effectively neutralizing HBV infection

The preparation of hybridoma producing mAbs against HBsAg resulted in establishment of nine hybridoma candidates, 7-4G, 11-4E-2H, 16-2G, 18-6B, 24-11D, 30-12G, 43-3D, 49-1B, and 66-2F. The neutralization activities for HBV infection of each mAb were evaluated using PXB cells and serum from a patient infected with HBV. The culture medium of each hybridoma was added at the indicated concentrations into the culture medium of PXB cells prior to infection with the HBV serum. Thirteen days after the inoculation, the production of HBsAg and HBeAg in the culture medium was measured by ELISA. As shown in Fig. 1a, and 1b, the monoclonal antibodies of seven clones, 7-4G, 18-6B, 24-11D, 30-12G, 43-3D, 49-1B, and 66-2F, showed suppressive effects on HBV infection but those of two clones, 11-4E-2H and 16-2G, did not. Comparing the effective concentration of the mAbs among the seven clones, the mAb from 7-4G clone, 7-4G mAb, was chosen to be analyzed further, as this antibody displayed the most effective neutralization at lower concentrations.

**Figure 1.**
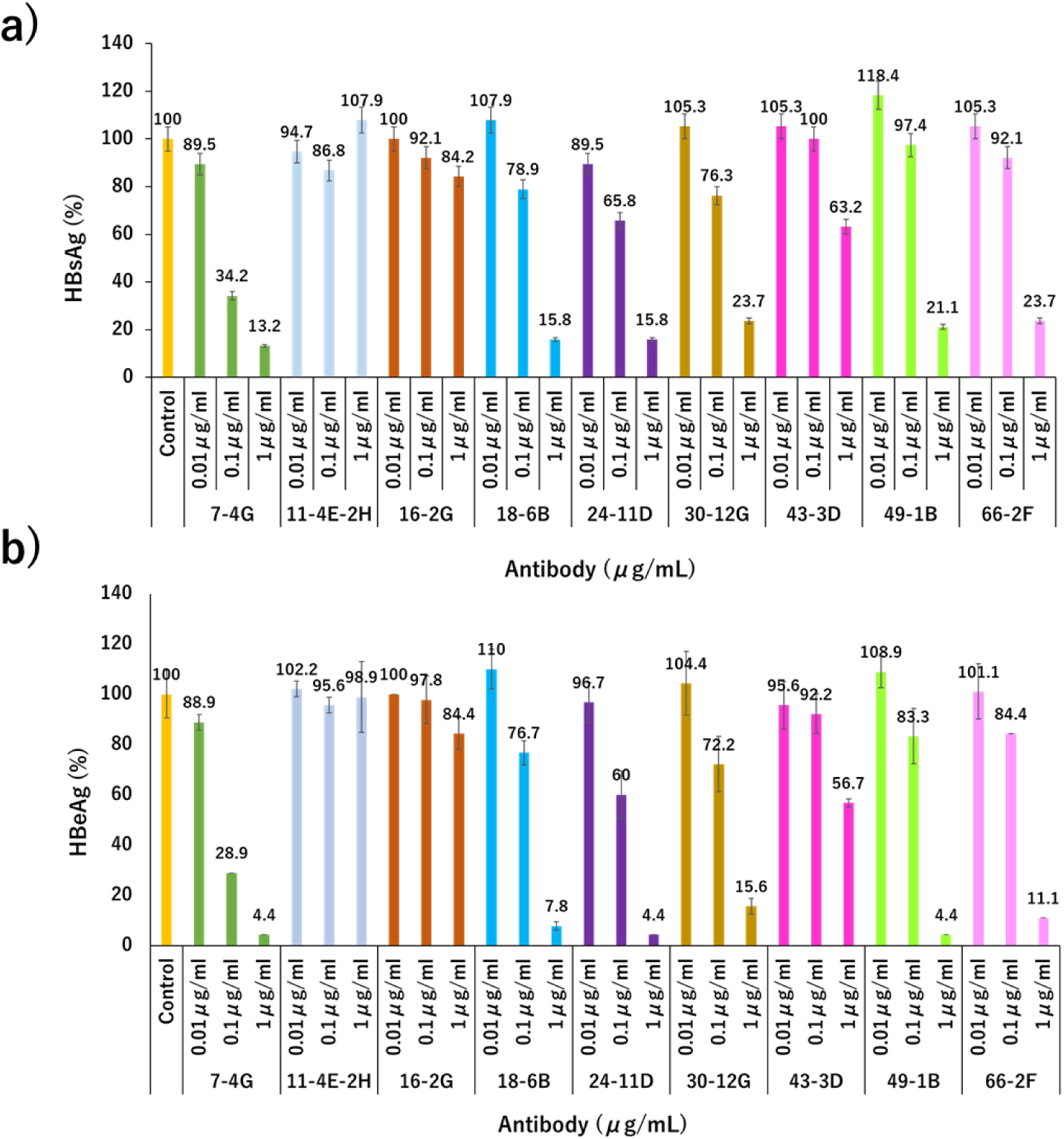
Neutralization activity of each selected mAb assessed in vitro. The culture supernatants of nine different antibody-producing hybridomas (mAbs at 0.01, 0.1, and 1 mg/mL) were added to PXB cells followed by HBV infection using serum from a patient infected with HBV (high HBV DNA copies/mL). After 24 hours, the cells were washed with PBS and cultured in fresh medium, which was exchanged every three or four days. After thirteen days, the amounts of HBsAg ***a)*** and HBeAg ***b)*** in the culture supernatants were measured by ELISA. The vertical axis shows the relative (percentage) values compared with results of the no antibody addition control, which is set at 100%. Data is presented as means ± standard errors of the means (SEM), and n=3.

### 2. Characteristics of the 7-4G mAb

To assess the antigenic structure of HBsAg recognized by the 7-4G mAb, at first, western blot analysis using 7-4G mAb following SDS-PAGE separating HBV fraction of patient’s serum was conducted. No polypeptide recognized by the 7-4G mAb, however, was detected (data not shown). The binding activity of the antibody, therefore, against the native form of HBsAg was examined by ELISA at first. As shown in Fig. 2a (blue line), 7-4G mAb exhibited a strong binding affinity to non-denatured HBsAg in an antibody concentration dependent manner. In contrast, the antibody showed minimal binding to HBsAg denatured by SDS and 2ME (Fig. 2a, orange line), highlighting its high specificity for conformational epitopes. Next, the association of 7-4G mAb with the non-denatured form of HBsAg contained in HBV particles isolated from the patient’s serum with ultracentrifugation was examined by immunoprecipitation. The protein contents of the HBV particles were visualized by CBB staining (Fig. 2b, left panel). The red and green arrows are estimated to be the p27 and p21 of small HBsAg, respectively, although polypeptides with higher molecular weights were not identified, possibly due of its extreme low amount. The immuno-reactive proteins with 7-4G mAb were then precipitated and analyzed by SDS-PAGE followed by silver staining (Fig. 2b, right panel, lane 2). The negative control experiment was similarly done by using normal mouse IgG (Fig. 2b, right panel, lane 1). Components that appear to be p27 and p21, indicated by the red and green arrows, respectively, were observed in lane 2 but not lane 1, confirming that 7-4G mAb recognized the native small HBsAg in the HBV related particles.

**Figure 2.**
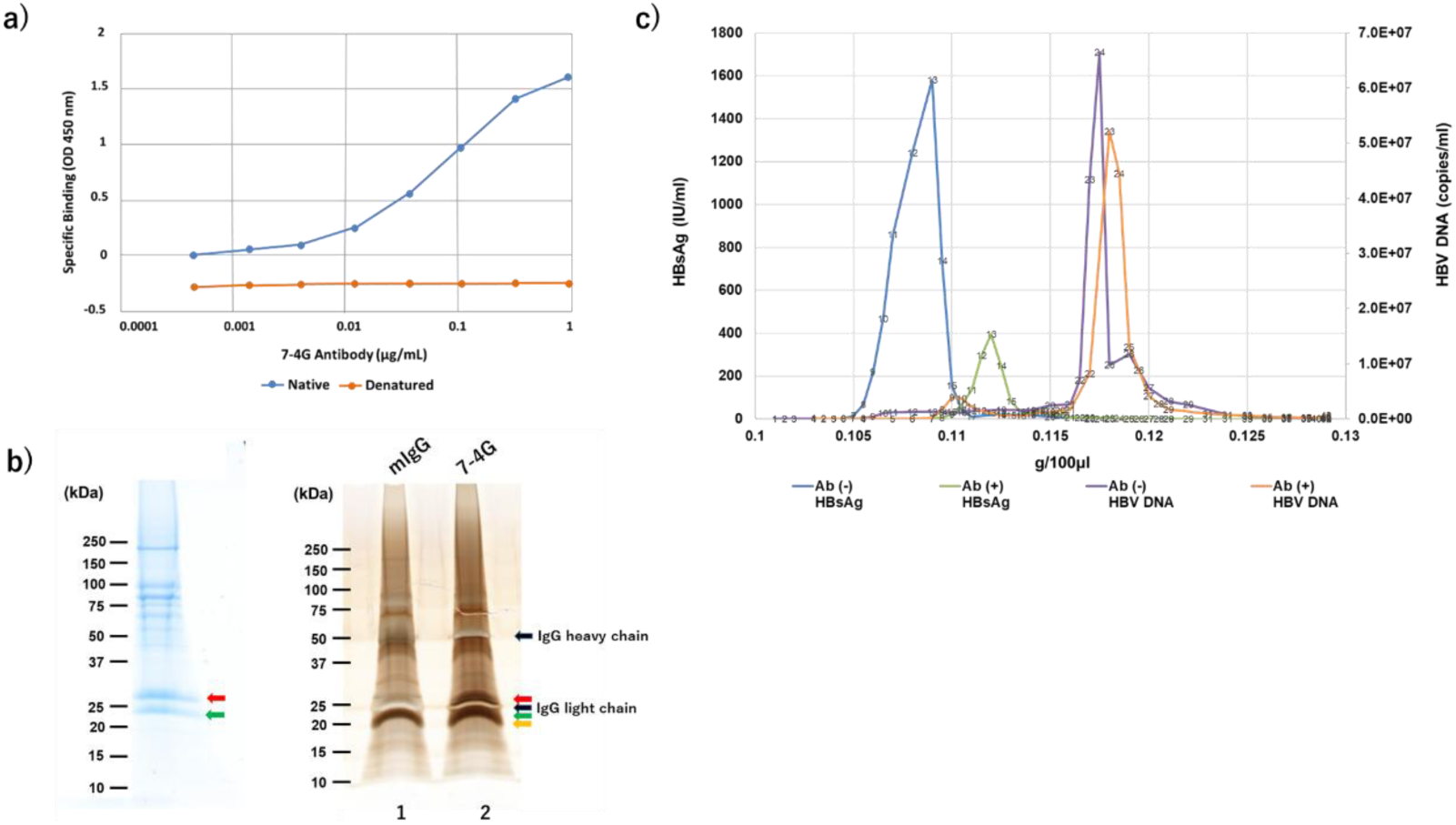
Characterization of 7-4G mAb. a) Structural property of HBsAg for binding with 7-4G mAb. The graph shows the results of ELISA indicating specific binding as an optical density (OD) of 450 nm (vertical axis) at different antibody concentrations (abscissa axis). The blue line and orange line shows the data of native and denatured HBsAg, respectively. ***b) Detection of the polypeptide immunoreactive with 7-4G in HBsAg from patient serum***. The HBsAg partially purified from patient serum was analyzed by SDS-PAGE followed by CBB staining (left panel). The immuno-reactive protein in the partially purified HBsAg was precipitated with 7-4G mAb (lane 2) or mouse IgG as negative control (lane 1) and analyzed by SDS-PAGE followed by silver staining (right panel). The red and green arrows show p27 and p21, the small HBsAgs, respectively. The black arrows show the unstained bands of IgG heavy and light chains. The yellow arrow indicates the probable nonspecific bands reactive against protein A Sepharose. ***c) Effect of 7-4G mAb on the buoyant density of HBV particles from patient serum.*** The results of buoyant density analysis, in which HBV particles from the serum of the patient with and without addition of 7-4G were separated by ultra-centrifugation and fractionated. The amounts of HBsAg and HBV DNA, indicated on the left and right vertical axes, respectively, in each fraction, of which buoyant density is from 0.1 to 0.13 g/100 μL as shown on abscissa, were plotted. The peak colored with green and blue show the results of HBsAg contents of the sera with and without 7-4G addition, respectively. The peak colored with orange and purple show the results of HBV DNA contents of the sera with and without 7-4G addition, respectively.

Furthermore, the HBV related particles recognized by 7-4G mAb in the serum from the patients infected with HBV were examined by fractionation. After addition of 7-4G mAb or PBS as a control, the serum was fractionated by buoyant density-gradient ultra-centrifugation and then the amounts of HBsAg and HBV DNA in each fraction were assessed. The peaks colored with orange and purple show the results of HBV DNA contents in the sera with and without 7-4G addition, respectively, indicating these fractions in the peaks include Dane particles (Fig. 2c). The purple peak shifted to the orange peak with heavier buoyant density in the presence of 7-4G mAb, probably because of binding of 7-4G mAb to the Dane particles. It should be noted that the total amount of DNA in the orange peak slightly decreased from that in the purple peak, likely due to the binding with 7-4G mAb. The peaks colored with green and blue show the results of HBsAg contents in the sera with and without 7-4G addition, respectively. (Fig. 2c). As these peaks do not contain HBV DNA, this denotes that the fractions constitute subviral particles (SVPs) mainly composed of small HBsAg. The blue peak shifted to the green peak with heavier density in the presence of 7-4G mAb, suggesting the binding of 7-4G mAb to SVPs. Prior results indicated that the target of 7-4G mAb is the native form of small HBsAg present on both HBV related particles. In addition, a particularly striking result was that the amount of HBsAg in the fractions of the blue peak was largely diminished in that of the green peak, probably showing that the HBV related particles including SVPs were reduced by binding of 7-4G mAb.

### 3. The inhibitory effect of 7-4G mAb on HBV infection in vivo

The efficacy of 7-4G mAb against HBV infection was also evaluated by an in vivo neutralization assay. Fig. 3a shows the schematic representation of the schedule for assessment of the neutralization activity of 7-4G mAb. We previously reported the activity of 3-10 mAb, which is included in the current study for synergistic comparison, as an antibody targeting specific regions within the preS1 domain of HBV genotype C (22). Five groups of mice, three mice in each group, were used for this study, whereby the antibodies were administered a day before HBV injection, and HBV DNA levels monitored weekly for eleven weeks. In the negative control group, where no antibody was injected, the levels of HBV DNA increased in the mice sera from the first week to the eleventh week after HBV inoculation (Fig. 3b). An almost similar pattern was observed in the second group, where the animals received the anti-FLAG antibody as a negative control (Fig. 3c). Although the group administered with 3-10 mAb showed a slight increase of HBV DNA after 5 weeks (Fig. 3d), in the fourth and fifth groups, where the mice received 7-4G mAb alone (Fig. 3e), or a combination of the two antibodies (7-4G and 3-10 mAbs), (Fig. 3f), respectively, none of the animals showed detectable levels of HBV DNA. These results suggested that the mAbs possess the potent neutralizing activity against the infection of HBV in vivo. In order to investigate the genotype dependency of the neutralizing activity of 7-4G mAb, three PXB mice per group were pretreated with the antibody followed by infection with HBV infected mouse sera of the other three most common HBV genotypes, besides genotype C namely; A, B, and D using the same protocol shown in Fig. 3a. HBV DNA levels were monitored weekly for eleven weeks. As demonstrated in Fig. 3b and 3c, mice untreated with antibody or treated with a control antibody develop HBV infection (genotype C), and HBV DNA is detectable and keeps increasing from the first week post-infection. However, pre-treatment with 7-4G mAb prevented the infection (Fig. 3e). This time, similar observations were made when 7-4G mAb-pretreated mice were infected with other HBV genotypes (A, B and D) as HBV DNA remained below detection limits for the same observation period of eleven weeks (Fig. 3g-3i). These data indicate that the neutralizing effect of the 7-4G mAb is independent of HBV genotypes examined so far.

**Figure 3.**
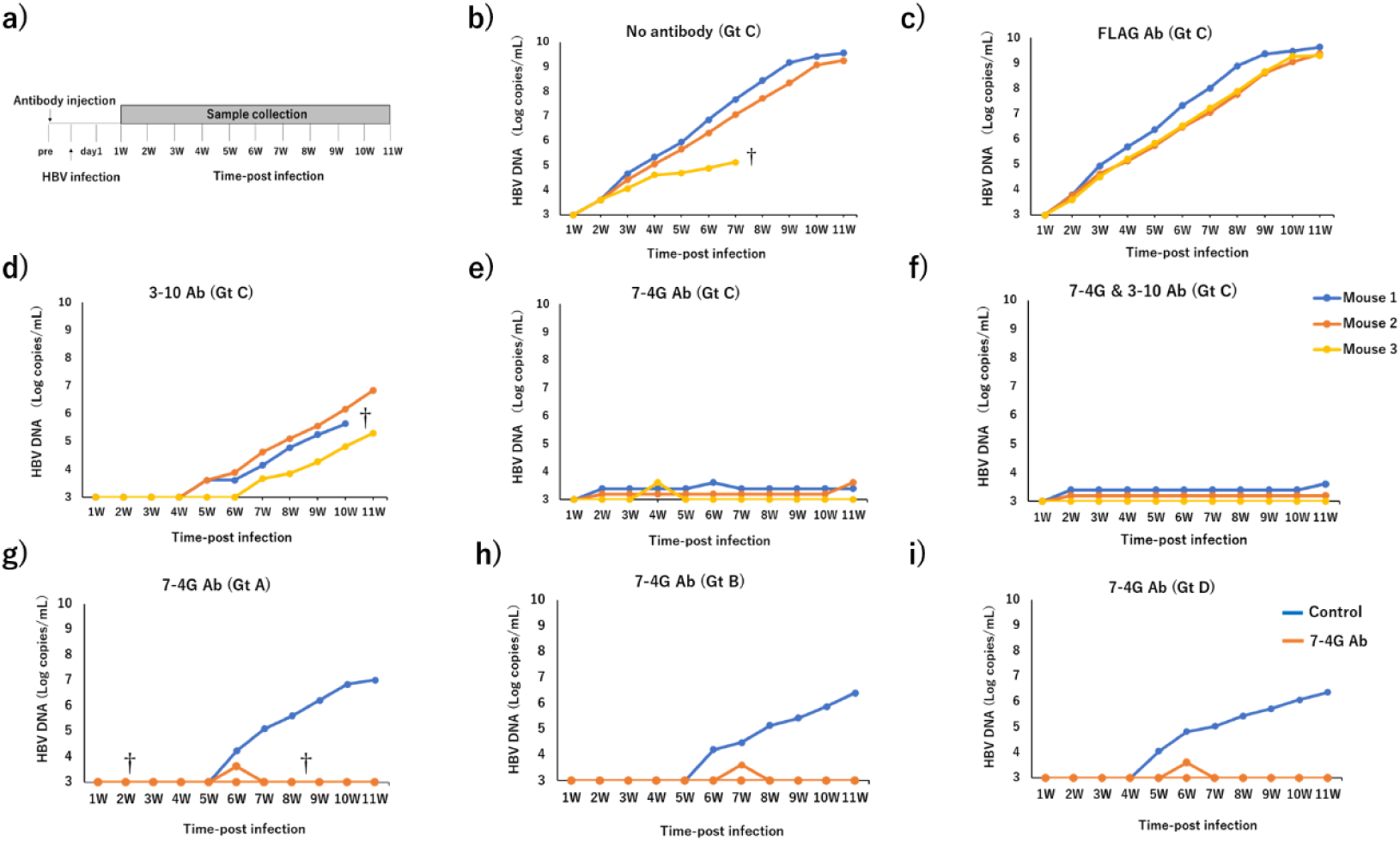
The inhibitory effect of 7-4G mAb Ab on HBV infection in vivo. *a)* Schematic representation of the schedule for assessment of the neutralization activity of 7-4G mAb using PXB mice in vivo. ***b-f)*** Three mice per group were pretreated with or without the antibody (ies) 24 hours before infection using a patient serum containing 10^3^ GEq HBV genotype C. The levels of HBV DNA in the sera at the designated time points (abscissa axis) were monitored and quantified by qPCR as shown on the left vertical axis as log-fold change to baseline. The antibodies used for pretreatment were 3-10 mAb (1 mg/mouse) (d), 7-4G mAb (0.5 mg/mouse) (e), 7-4G mAb (0.33 mg/mouse) in combination with 3-10 mAb (0.66 mg/mouse) (f). The anti-FLAG Ab (0.5 mg/mouse) was used as a negative control (c) as well as no antibody administration (b). ***g-i)*** The inhibitory effects of the 7-4G mAb on other HBV genotypes, namely, genotype A (g), genotype B (h), and genotype D (i) were similarly assessed. However, in experiment g-i, the PXB mice were injected with 10^3^ GEq HBV sera derived from mice chronically infected with the respective HBV genotype, subsequent to 24-hour pre-administration of 50 mg/kg/mouse of 7-4G antibody. In b-f, each color represents one mouse, and for g-i, the blue color indicates the control antibody treated mouse, while the orange color indicates mice treated with the 7-4G mAb. † indicates death of a mouse.

### 4. The effect of mAb treatment on chronic HBV infection in mouse model

The effects of the mAbs on chronic infection of HBV were examined next using PXB mice inoculated with HBV. The mice sera were monitored from two weeks after the inoculation of HBV to monitor HBV DNA levels, which showed active replication and production of HBV (data not shown). HBV DNA started to gradually increase in all mice from the second week, running to peak at around the 15th week, at which the antibodies were administered. Polygonal lines indicating the HBV DNA levels in the control (FLAG antibody administered) mice kept the highest plateau levels for the entire 8 weeks of observation (Fig. 4a).

**Figure 4.**
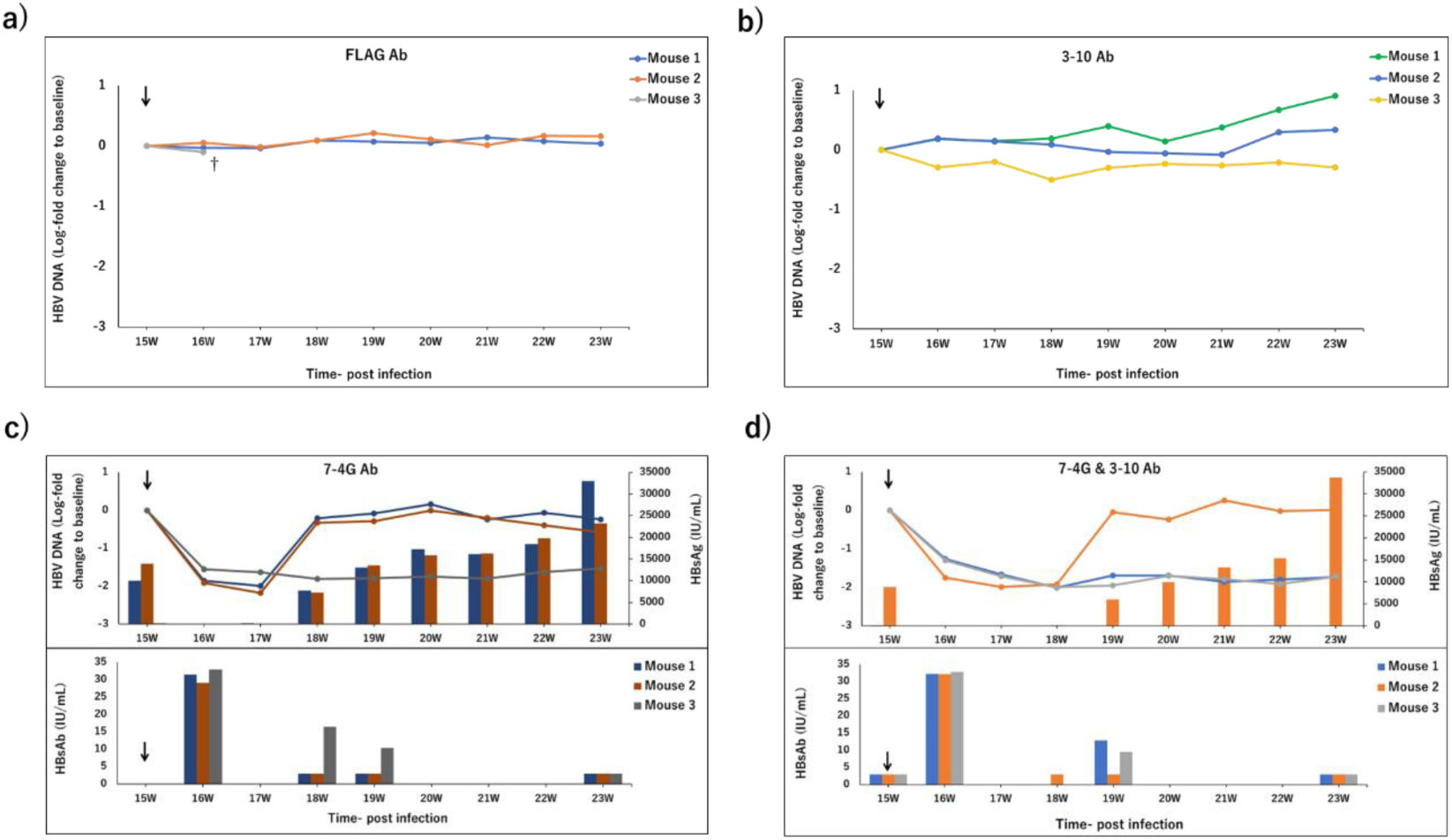
The effect of 7-4G mAb on chronic HBV infection. PXB mice chronically infected with HBV genotype C were established by inoculation with patient serum and monitored for HBV DNA levels from two weeks post infection. After confirmation of the attainment to the plateau level of HBV DNA in the mice sera at fifteen weeks post infection, the injection of the antibodies was started (indicated by arrows). The levels of HBV DNA in the sera at the designated time points (abscissa axis) were monitored for eight weeks, and quantified by qPCR as shown on the left vertical axis (log-fold change to baseline). Three mice were assigned per group. ***a)*** HBV DNA levels at the designated time points after inoculation in the sera from mice with anti-FLAG mAb, and ***b)*** with 3-10 mAb antibody treatments, are shown. ***c)*** The data of HBV DNA levels of experimental groups, in which 1 mg/mouse of 7-4G mAb alone, or ***d)*** 1 mg/mouse of 7-4G mAb in combination with 2 mg/mouse of the 3-10 mAb were injected, are shown, respectively. In addition to the HBV DNA data in c and d, HBsAg levels (IU/mL) in the sera (right vertical axis, upper panels), as well as the levels of HBsAb (IU/mL) (left vertical axis, lower panels) at those time points were also measured. In a-d, each color represents one mouse. † indicates death of a mouse.

In the 3-10 mAb-treated group, HBV DNA levels in the sera of those mice also kept the plateau levels following injection with the mAb (Fig. 4b, blue and green lines), although in one mouse, HBV DNA slightly decreased for three weeks after antibody injection but then returned to the highest plateau levels for the remainder of the study period (Fig. 4b, yellow line).-These data showed that the proliferation and chronic infection of HBV in the mice were not largely affected by the injection of 3-10 mAb, at least in this condition.

In the 7-4G mAb-treated group, HBV DNA levels in the sera of all three mice showed the rapid and double-digit decrease for two weeks after the single 7-4G mAb injection, where this time is the restrain stage of the treatment (Fig. 4c, mazarine, red and gray lines in upper panel). Thereafter, HBV DNA levels in two of the three mice returned to the highest plateau levels three weeks after the injection, indicating that HBV in the mice started to proliferate and to produce HBV particles, (Fig. 4c, mazarine and red lines in upper panel), while the decreased HBV DNA level was sustained for eight weeks after the injection in the serum of one mouse (Fig. 4c, gray line in upper panel). HBsAg levels in the sera of two mice also showed the decrease to the marginal level until two weeks after the injection. This however, returned to detectable levels, followed by a gradual increase until eight weeks post-injection (Fig. 4c, mazarine and red bars in upper panel). It is noteworthy that HBsAg levels of the other one mouse was barely detected in the entire experimental period (Fig. 4c, gray bars in upper panel). HBsAb was detectable in the sera of all three mice one week after the injection of antibody at a high level (30 IU/mL). Thereafter, HBsAb levels in the sera of two mice decreased rapidly to low levels (3 IU/mL) (Fig. 4c, mazarine and red bars in lower panel), while that in one mouse showed a gradual decrease from three to eight weeks after the injection (Fig. 4c, gray bars in lower panel). These results showed that injection with 7-4G mAb caused the transient decrease of HBV DNA and HBsAg levels in the sera of mice chronically infected with HBV. And remarkably, a prolonged decrease of HBV DNA was observed in a mouse with a relatively long presence of the injected mAb in the serum.

In the group treated with 7-4G mAb coupled with 3-10 mAb, HBV DNA levels in the sera of all three mice showed a rapid and double-digit decrease for three weeks after mAbs injection, at the restrain stage of the treatment (Fig. 4d, blue, orange and gray lines in upper panel). Thereafter, the decreased HBV DNA level was sustained for eight weeks after the injection in the sera of two mice (Fig. 4d, blue and gray lines in upper panel), although HBV DNA levels in one mouse returned to the highest plateau levels four weeks after antibody injection (Fig. 4d, orange line in upper panel). HBsAg in the serum of one mouse also showed the decrease to the marginal level until three weeks after the mAbs injection, then returned to be detected, followed by a gradual increase until eight weeks post-injection (Fig. 4d, orange bar in upper panel), while HBsAg in the sera of the other two mice were barely detectable during the entire experimental period (Fig. 4d, blue and gray bars in upper panel). HBsAb was observed in the sera of all three mice one week after the injection of antibody at high level. Those in the sera of one mouse decreased rapidly to low levels (Fig. 4d, orange bars in lower panel), while those in two mice showed a gradual decrease from four to eight weeks after injection (Fig. 4C, blue and gray bars in lower panel). These results showed that the co-administration also caused the transient decrease of HBV DNA levels in the sera of mice chronically infected with HBV. And the prolonged decrease of HBV DNA was observed in two of three mice with relatively long presence of injected mAb in the sera, just as in the case of 7-4G mAb single injection. These data showed that, despite the presence of the antibody as much as little in the serum, the proliferation and chronic infection of HBV in the mice were not largely affected by the injection of 3-10 mAb.

### 5. Combination treatment with ETV and 7-4G mAb in mouse model

ETV is one of the NRTIs, anti-HBV drugs which work through inhibition of viral replication (4). As the potent anti-HBV activities of 7-4G-mAb were observed in PXB mice chronically infected with HBV, the effects of the mAb in combination with ETV were examined next. PXB mice inoculated with HBV as above were either daily administrated with ETV from the ninth (peak of HBV infection) to the eighteenth week after inoculation (ETV alone group, blue lines), or in addition to ETV treatment, injected with 7-4G mAb at the thirteenth and sixteenth weeks after inoculation (ETV and 7-4G group, orange lines). As shown in Fig. 5, the high levels of HBV DNA observed in the sera of all six mice at the ninth week after the inoculation started to decrease in double and single digits one and two weeks, respectively after the administration of ETV. Thereafter, there was a relatively gradual decrease for five weeks, ending in a plateau from about week sixteen for the ETV alone group (Fig. 5, mazarine lines). On the other hand, the injection of 7-4G mAb at week thirteen caused a further double digits decrease of the HBV DNA level in the serum of one mouse and single digits decrease in the other two mice (Fig. 5, orange lines) in one week. One mouse died at week sixteen, but in the remaining two mice, re-injection with 7-4G mAb at this time point caused a further decrease in HBV DNA levels showing that the 7-4G mAb injection after ETV treatment has an additive suppressive effect on the chronic HBV infection of the mouse model.

**Figure 5.**
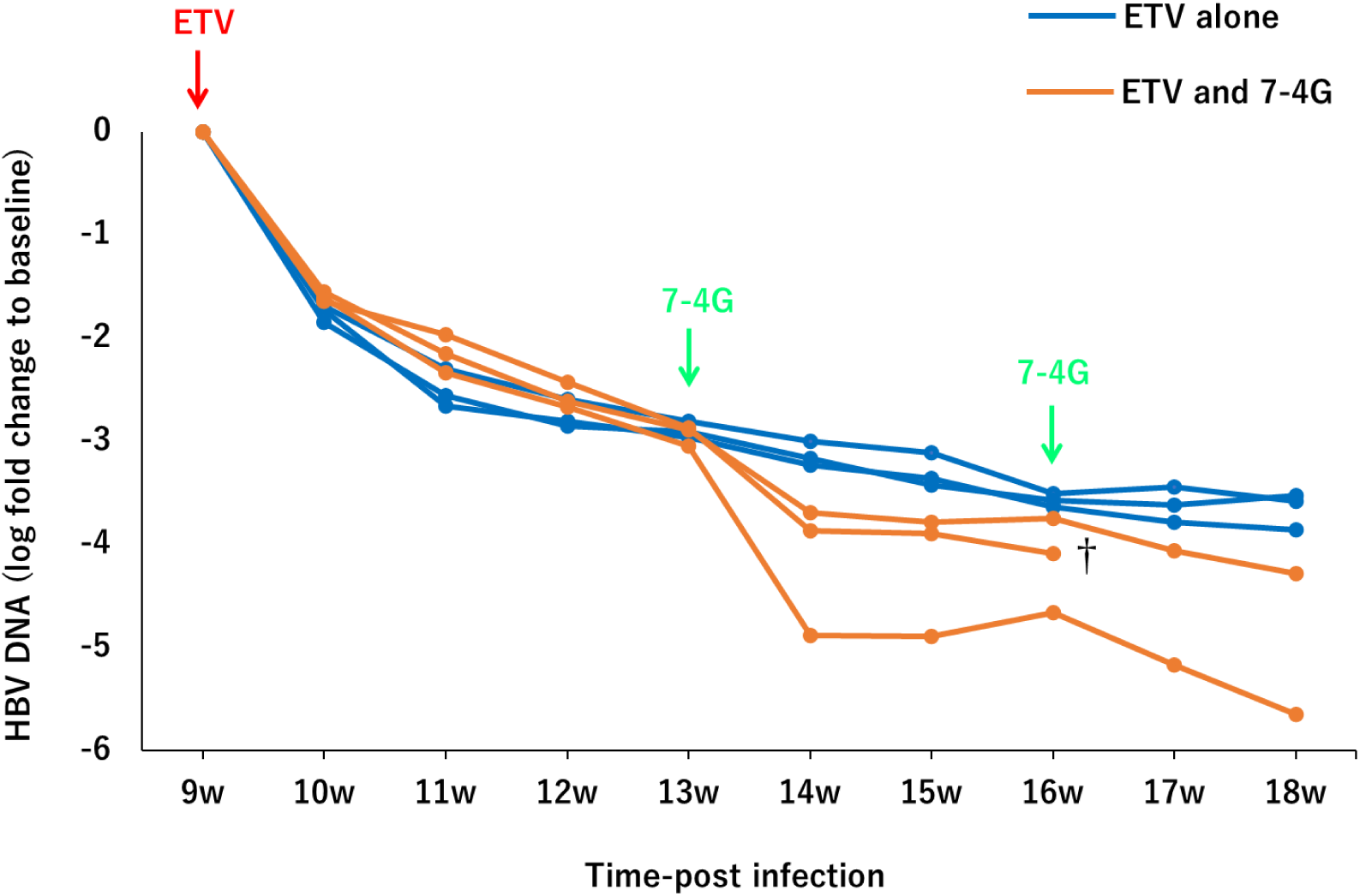
The effect of combined administration of ETV and 7-4G mAb. Administration of ETV (0.2 mg/kg/day) to six PXB mice in a chronically HBV infected state established nine weeks after HBV inoculation was started, as indicated by the red arrow, and continued to the end of the study at eighteen weeks post infection. The injection with (orange lines) or without 1 mg/mouse of 7-4G mAb (blue lines) in the mice was initiated at the thirteenth-and sixteenth-week post HBV inoculation, as shown by the green arrows. The mice sera were collected every week from the ninth week (the start of ETV treatment) till the end of the study period at eighteen weeks post HBV inoculation. The levels of HBV DNA in the sera at the designated time points (abscissa axis) were quantified by qPCR and shown on the left vertical axis as log-fold change to baseline. † indicates death of a mouse.

## Discussion

In the current study, a novel monoclonal antibody, 7-4G mAb, targeting the HBsAg was developed and characterized. The antibody was demonstrated to recognize the conformational epitope on the native form of the glycosylated small HBsAg, p27 (Fig. 2) and to possess a neutralization activity against HBV infection both in vitro and in vivo (humanized liver chimeric mouse model) (Fig. 1 and Fig. 3, respectively). There have been many reports of such mAbs derived from human as well as mouse (14,23), preventing HBV infection without direct association with the pre-S1 region of the large HBsAg of which interaction with the HBV receptor molecule, sodium taurocholate co-transporting polypeptide (NTCP) is known to be essential for HBV infection (24). For example, those mAbs binding may induce the conformational change of large HBsAg through the interaction with its small HBsAg part or cause the spatial hindrance, resulting the disruption of the interactions between HBsAg and cell surface proteins, such as NTCP and the low-affinity cell receptor heparan sulfate proteoglycan (HSPG) (25,26).

It is notable, however, that the reduction of HBV related particles, especially SVPs, in the patient serum caused by 7-4G mAb was observed in vitro (Fig. 2c). It seems likely that the HBV related particles including HBsAg were disrupted by 7-4G mAb in vitro, probably via complement-dependent virion lysis (virolysis) using complement present in sera from the patients infected with HBV (27–29). This observation is likely to imply that the 7-4G mAb-dependent neutralization of HBV infection observed in vitro and in vivo was due to its potential reduction of HBV related particles in addition to the neutralization activity described above. The rapid and double-digit decrease of HBV DNA by the single injection of 7-4G mAb within one week observed in PXB mice chronically infected with HBV also seemed to be due to, at least partly, this virolysis potential of 7-4G mAb. On the other hand, treatment with Myrcludex B, an HBV entry inhibitor composed of a preS1 polypeptide suppressing interaction of preS1 of HBsAg with NTCP, in PXB mice with chronic HBV infection did not show such rapid and strong reduction of HBV DNA (30). It is, therefore, unlikely that only the neutralizing activity causes such a rapid HBV particle reduction. As PXB mouse model was developed from severe combined immunodeficient mouse (SCID), the acquired immune responses are not present in the mouse (19). The complement system, as well as innate immune system, however, still exists in the mouse because human hepatocytes producing the complement were engrafted in the mouse (19). Therefore, the rapid decline of the HBV related particles in the PXB mice treated with single shot of the mAb(s) seemed to show the possible contribution of the complement dependent system in the mice, including complement-dependent virolysis and phagocytosis of opsonized viral particles (31). The precise mechanism of HBV particle reduction by 7-4G mAb should be examined further both in vitro and in vivo, because the anti-preS1 mAb, 3-10 mAb, did not show such an effect on the HBV DNA present in the sera from PXB mice chronically infected with HBV (Fig. 4b).

One of the remarkable results of this study is the long-lasting suppressive effects of the single shot treatment of 7-4G mAb on markers of chronic HBV infection in the mouse model (Fig. 4c and 4d). HBV particle production in sera remained suppressed for the entire eight weeks of observation in one of three mice treated with 7-4G mAb alone and in two of three mice in the anti-preS1 co-administered case. The molecular mechanism of this sustained suppression of HBV markers in the sera of PXB mice remains unclear yet. However, it seems possible that 7-4G mAb purges the HBV infected hepatocytes in the PXB mice by complement-dependent cytotoxicity through the membrane attack complex formation, just like as probable complement-dependent virolysis by 7-4G mAb as described above. Of course, the antibody dependent cellular cytotoxicity (ADCC) might play a role in elimination of HBV infected hepatocytes in the PXB mice as well, because natural killer (NK) cell-function of ADCC is present in the SCID mouse (27,31,32). Otherwise, the internalized 7-4G mAb in the HBV infected hepatocytes may inhibit the egression of HBV particles as described previously (33,34). It seems credible that the sustention of the antibody is one of the determinants for the effect of treatment, as the relatively prolonged presence of 7-4G mAb in sera was observed in all the mice that showed the long-term suppressive effects (Fig. 4c and 4d). This may be supported by the additive effect of 3-10 mAb, the anti-preS1 antibody, whose combination with 7-4G mAb showed prolonged suppression of HBV DNA in the early phase of the treatment (Fig. 4d), in spite of no suppressive effect by 3-10 mAb treatment alone (Fig. 4b). The dose-dependent response of the effect, therefore, should be examined in the future. On the other hand, the multi-epitope approach, may be consistent with the previous reports showing that cocktails of antibodies directed against distinct HBV surface domains yield broader and more durable antiviral activity than single antibody alone (35). Considering the use for treatment after humanization of the mAbs (36), concurrent targeting of small HBsAg and preS1 epitopes may enhance immune-mediated clearance of infected hepatocytes through Fc-dependent pathways, and contribute to sustained viral control even after antibody levels decline. Further analysis with an enlarged sample size of the animal experiments and various treatment conditions is required to clarify the precise molecular mechanisms of the anti-HBV effects of 7-4G mAb.

Furthermore, co-administration of 7-4G mAb with entecavir resulted in even greater suppression (Fig. 5), indicating the potential use of the antibody in combination therapy for improved HBV treatment outcomes. NAs such as entecavir and tenofovir remain first-line agents for chronic HBV infection for preventing a cancer development (37). These drugs effectively inhibit reverse transcription and suppress HBV DNA replication to undetectable levels in most patients. However, NAs cannot eliminate the cccDNA of HBV or integrated HBV DNA, leading to viral rebound when treatment stops. Furthermore, the production of viral gene products, which were suggested to be related to cancer development, continued from those HBV DNAs during the course of medication (38). In the present study, entecavir monotherapy produced a rapid, double-digit decrease in HBV DNA one week after administration. The subsequent addition of 7-4G mAb caused a further two or one-log reduction in HBV DNA, reaching low single-digit levels by week thirteen (Fig. 5). This 7-4G mAb-dependent reduction in HBV DNA is likely due to decrease of the circulating HBV-related particles including Dane particles (Fig. 2c) and possibly HBsAg producing cells that NAs alone cannot achieve (39).

Monoclonal antibody therapy offers several practical advantages. Antibodies are highly specific, generally safe, and can be engineered for extended half-life or enhanced effector function (40). Compared with other emerging HBV treatments such as therapeutic vaccines, RNA-silencing agents, or entry inhibitors, antibody-based therapy holds unique strengths (40). Vaccines and RNA drugs can reduce viral markers but depend on variable host immune responses, while entry inhibitors block only new infections. Antibodies combine both entry blockade and antigen removal as shown in this study, while simultaneously engaging immune effector pathways (41). Future investigations may aim to extend antibody duration and potency. For instance, Fc-engineering to increase FcRn binding or generation of bispecific antibodies that target multiple HBV epitopes may improve therapeutic outcomes (42,43). Moreover, combining antibody therapy with immune modulators such as checkpoint inhibitors or therapeutic vaccines could also enhance viral clearance (44). Collectively, these efforts will help define the role of antibody therapy in achieving sustained functional cure.

In conclusion, the monoclonal antibody 7-4G displayed strong and specific antiviral activity against HBV in both cell culture and humanized mice models. Its combination with entecavir produced deeper viral suppression than either treatment alone. These findings highlight antibody-based therapy as a promising addition to current HBV treatment strategies and a potential component of future combination regimens aimed at a functional cure for chronic hepatitis B.

## Acknowledgement

This study was funded by the Program for Basic and Clinical Research on Hepatitis of the Japan Agency for Medical research and Development (grant number: JP25fk0310532h0001).

